# Cost adjusted hierarchical defense strategies enables stratified oxidative stress tolerance

**DOI:** 10.64898/2026.03.03.709455

**Authors:** Anjali V. Patil, Jiao Zhao, Snehal V. Khairnar, Sonali Srivastava, Laurence Yang, Amitesh Anand

**Affiliations:** Department of Biological Sciences, Tata Institute of Fundamental Research, Mumbai, Maharashtra, India; Department of Chemical Engineering, Queen’s University, Kingston, Ontario, Canada; BRIC-National Institute of Immunology, Delhi, India

## Abstract

Organisms employ diverse adaptive strategies to withstand environmental stresses, yet how bacteria tailor their responses to different magnitudes of the same stressor remains poorly understood. This question is particularly salient for oxidative stress, which varies substantially across physiological niches and exerts distinct selection pressures on colonizing bacterial populations. Here, we used four separate adaptive laboratory evolutions of *Escherichia coli* across multiple paraquat concentrations and genetic backgrounds to dissect how cells adapt to varying levels of superoxide stress. Integrating multi-omic analyses with tailored genome-scale metabolic modeling, we identify two fundamentally distinct tolerance strategies based on flux control and cellular resource optimization. Under low superoxide stress, blocking polyamine transporter that is hijacked for paraquat influx suffices to maintain redox balance. In contrast, higher stress levels activate an energetically demanding program involving enhanced detoxification of reactive radicals and active efflux that allows cells to maintain higher aerotype. We further demonstrate that flux regulation establishes a primary defense layer, upon which metabolic repair systems act to provide additional fitness advantages. Together, our findings reveal how *E. coli* differentially engages modular stress-response programs depending on stress magnitude, offering generalizable insights into the principles governing dynamic bacterial adaptation.

## Introduction

Stress mitigation is fundamental to life, and organisms have evolved diverse adaptive strategies to endure environmental perturbations ^1^. Bacteria are directly exposed to their environment, which is often dynamic and hostile. Bacterial pathogens, therefore, must survive a multitude of stresses like nutrient availability, pH, oxidative stress, and thermal stress, among other factors, to colonize various niches ^2^. Despite possessing a relatively simpler genome, bacterial stress response pathways are highly versatile in mitigating a wide array of stress conditions ^3–5^. While the bacterial responses to distinct stresses are relatively well characterized, the mechanisms underlying adaptation to varying magnitudes of a stress remain poorly understood. This gap is especially relevant because physiological niches, such as distinct host tissues, impose variable oxidative environments and thereby exert differential selection pressures on colonizing populations ^6,7^.

Oxidative stress, arising from reactive oxygen species (ROS), represents one of the most pervasive threats encountered by bacteria, arising from diverse sources such as host immune responses, metabolic disturbances, and antibiotic exposure ^8^. Bacteria are equipped with a suite of ROS defense pathways to orchestrate a tailored response to the specific nature of oxidative stress. The transcriptional regulators SoxR, OxyR, and RpoS can evoke specific transcriptional changes to restore redox homeostasis ^9^. Additionally, chronic exposure to oxidative stress can lead to genetically programmed metabolic changes ^10,11^. However, such genetically fixed adaptive changes often have a fitness trade-off associated with them ^11^. Therefore, a careful resource allocation for growth versus defense systems governs optimal adaptive physiology.

We performed four separate adaptive laboratory evolutions (ALE) of *Escherichia coli* under different paraquat concentrations and with varied genetic backgrounds to delineate adaptive strategies to distinct magnitudes of superoxide stress. Based on the multi-omic characterization of the evolved strains and tailored genome scale model based computations, we report two distinct oxidative stress tolerance strategies in *E. coli* that are differentially engaged depending on the level of superoxide exposure. At low concentrations, limiting intracellular paraquat influx is sufficient for survival, whereas higher levels trigger an energetically costly program involving ROS detoxification and active efflux. The flux control based stress mitigation strategies allow further tolerance development using metabolic alterations. These findings highlight how cells modulate adaptive programs in response to the magnitude of stress, with implications for understanding the principles of dynamic stress-responsive regulation.

## Results and discussion

### Adaptive response of *E. coli* to varying superoxide levels

Respiratory detoxification of oxygen alone is insufficient to prevent oxidative stress; consequently, bacteria rely on transcriptional programs to maintain redox homeostasis during transient fluctuations in ROS levels ^8,12,13^. However, sustained exposure to reactive radicals necessitates broader adaptive strategies, including lifestyle shifts and metabolic reprogramming ^8,10^. While there is a general understanding of multiple strategies associated with ROS tolerance, the mechanistic distinction and stress level specialization is lacking ^10^. To investigate mechanisms underlying long-term adaptations, we performed ALE of *E. coli* under two distinct paraquat concentrations differing by an order of magnitude (Figure 1A). Three independent lineages were evolved in each condition to derive common adaptive traits and ALE was continued until growth-rate improvements plateaued. Strains evolved at 10 µM paraquat are hereafter designated LPE-A/B/C (Low Paraquat Evolved), and those evolved at 100 µM paraquat are designated HPE-A/B/C (High Paraquat Evolved) (Supp. Table 1).

**Figure 1:**
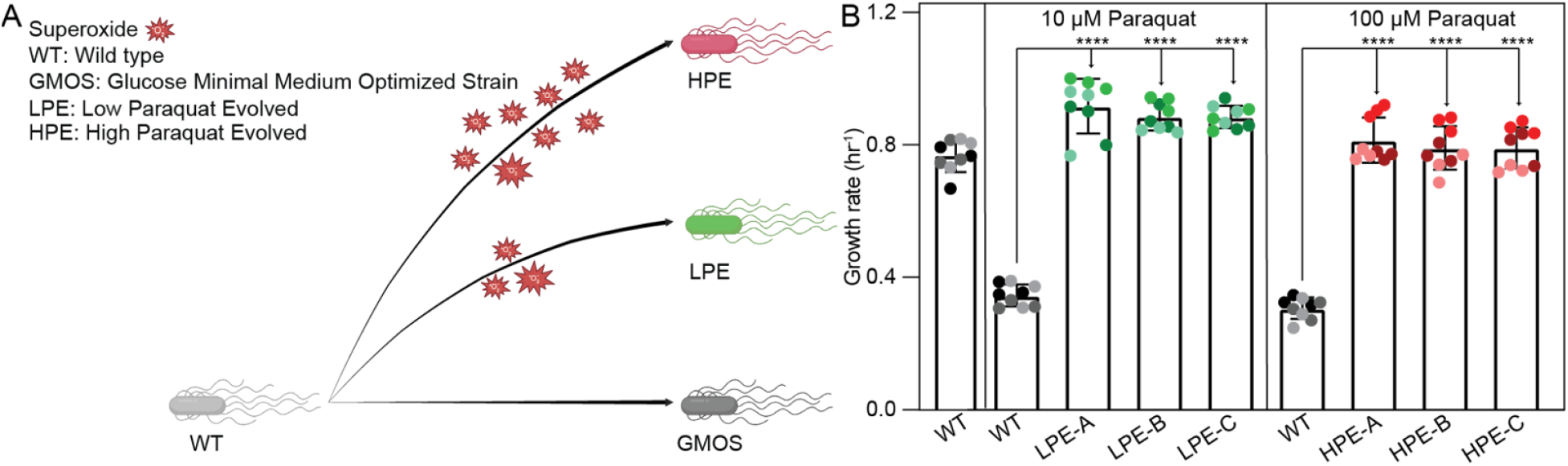
Adaptive evolution of *E. coli* in presence of different levels of superoxide. (A) Schematic of the adaptive laboratory evolution (ALE) design. (B) Growth rates of *E. coli* before and after ALE in the presence of 10 μM and 100 μM paraquat. The plot shows the mean of three biological replicates (each with three technical replicates (in the same color shade)), and the error bars show the standard deviation. Significance was determined using the Mann-Whitney test (****: P-value < 0.0001).

As expected, wild-type (WT) showed a significant growth rate drop at both paraquat concentrations (Figure 1B). Evolved strains achieved significantly higher superoxide tolerance and recovered their growth rate (Figure 1B). In paraquat free medium, LPE and HPE showed growth profiles similar to a WT strain, GMOS, that is evolved to grow to its maximum growth rate in M9-minimal medium with glucose as carbon source (Supp. Figure 1) ^14^. In the presence of paraquat, while GMOS fails to grow, LPE and HPE show robust growth. Interestingly, the LPE strains are specialised for lower paraquat tolerance, as their growth was limited at higher levels of paraquat with an extended lag phase (Supp. Figure 1). In contrast, HPE strains were tolerant to both paraquat concentrations. These phenotypic distinctions suggested a stress specialization and difference in underlying adaptive strategies between LPE and HPE.

### Distinct strategies to tolerate different magnitudes of superoxide stress

Stable adaptive phenotypic changes are often driven by genetic selection ^15^. We performed whole genome sequencing of the evolved strains to compare their genotypes. Both LPE and HPE acquired a common set of mutations that are well characterized to improve the growth of *E. coli* on M9 minimal medium (Supp. Table 2). The mutations in RNA polymerase subunits as well as in the intergenic region of *pyrE* and *rph* genes facilitates growth-supporting cellular resource allocation ^14–16^. This observation explains the growth similarity of LPE and HPE with GMOS in paraquat free medium (Supp. Figure 1). Remarkably, besides these common growth-promoting mutations, LPE and HPE strains showed mutations that were specific to stress levels.

All three independently evolved lineages of LPE acquired mutations in an ABC transporter PotD-PotABC. While this transporter is responsible for the uptake of polyamine spermidine, the structural analogy can enable cellular entry of paraquat through this system ^17–19^. LPE-A and LPE-B acquired mutations in *potC* and *potD*, respectively, that truncated the open reading frames (Figure 2A). In contrast, a point mutation in LPE-C results in PotA G55D substitution within its ATP-binding domain. This substitution introduces a negatively charged side chain and alters the residue’s helical propensity, which may disrupt protein folding ^20,21^. Such mutations should result in loss of function of this transporter and potentially limit paraquat influx. We generated a *potD* knockout strain to establish the role of PotD-PotABC disruption in paraquat tolerance. WTΔ*potD* displayed comparable growth to the WT strain under non-stress conditions but exhibited improved tolerance to 10 μM paraquat (Figure 2B). These results suggest that inactivation of the PotD-PotABC transporter confers growth advantage under paraquat stress by limiting the non-specific paraquat entry into the cell.

**Figure 2:**
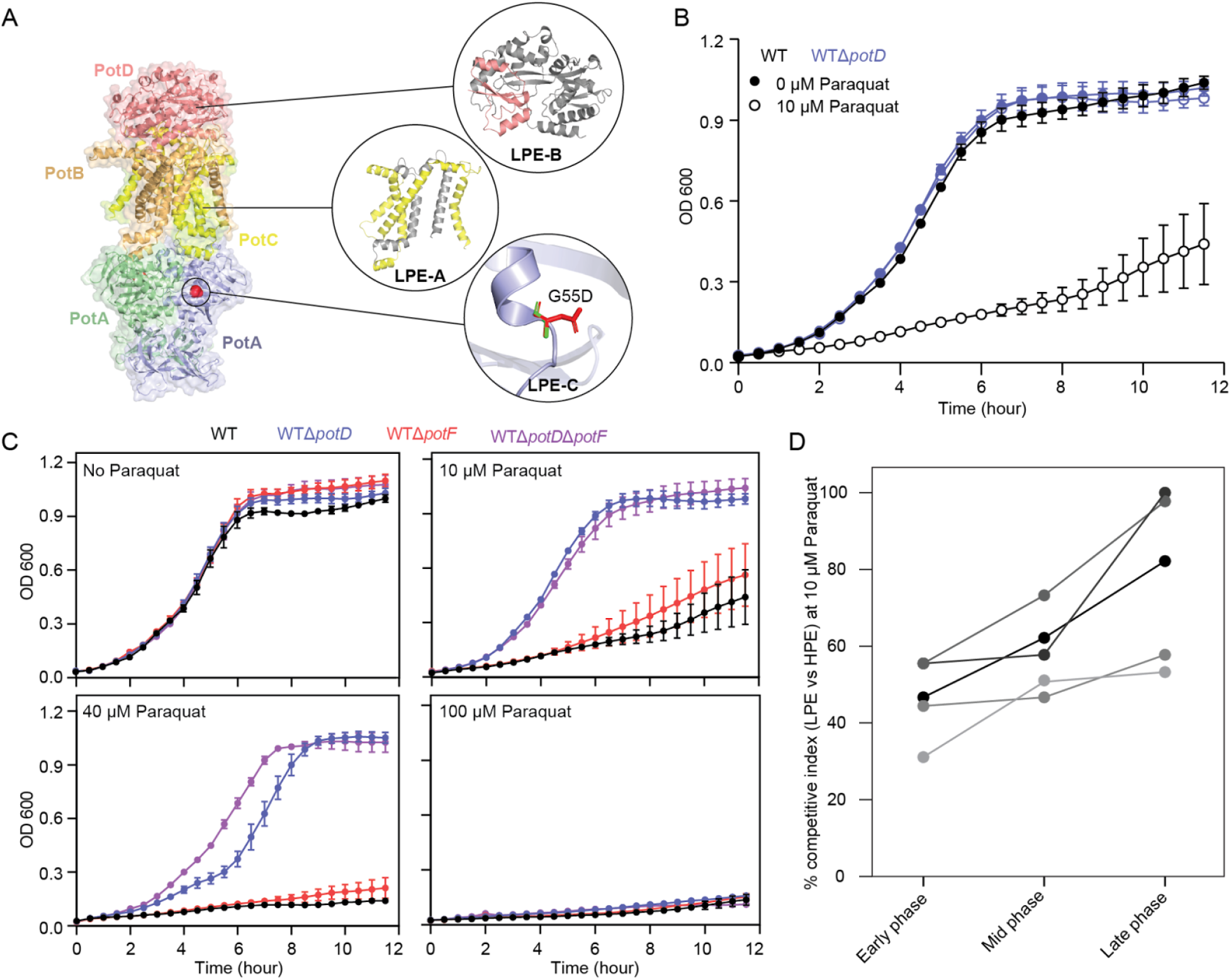
Adaptive strategy of oxidative stress tolerance in low-paraquat evolved *E. coli*. (A) The mutations observed in the LPE strains mapped on the PotD-PotABC complex from *E. coli* (Structure of PDB ID 8Y5I rendered using PyMol). In the zoomed-in insets of PotD and PotC, the regions shown in grey depict the truncated region. In the zoomed-in inset of PotA, green represents wild-type amino acid residue, while red represents mutant amino acid residue. (B) Effect of *potD* gene deletion on the growth profile of WT strain in the presence of 10 μM paraquat. The growth curve shows a mean of four biological replicates (with three technical replicates each), and the error bars show the standard error of the mean. (C) Effect of *potD* and *potF* gene deletion on the growth profile of WT strain in the presence of different paraquat concentrations. The growth curve shows a mean of four biological replicates (with three technical replicates each), and the error bars show the standard error of the mean. (D) Percent competitive index of LPE vs HPE in presence of 10μM paraquat. All five individual biological replicates shown on the plot.

*E. coli* also encodes a second polyamine uptake system, PotF-PotGHI, which is specific for putrescine ^22^; however, this pathway was not selected during growth optimization under paraquat stress. To examine the basis of this apparent evolutionary bias, we compared paraquat tolerance in WTΔ*potD* and WTΔ*potF* strains (Figure 2C). In contrast to *potD* deletion, loss of *potF* alone did not confer an appreciable tolerance at 10 μM paraquat. Notably, simultaneous deletion of *potF* and *potD* further enhanced tolerance relative to WTΔ*potD*. These results explain the preferential selection of the PotD-PotABC system and indicate that PotF-PotGHI does not serve as the primary route for paraquat influx. An important question emerging from this observation is whether the distinct substrate specificities for individual polyamine species influences their differential contribution to paraquat uptake ^17^. Notably, this import restriction based strategy provides limited tolerance and at higher paraquat concentrations the strains were significantly susceptible. However, the adaptive strategy of LPE seems to be better suited for lower paraquat concentrations than HPE. We performed competitive growth profiling by co-inoculating LPE and HPE in a medium containing 10 μM paraquat. We observed a progressive increase in the representation of LPE cells over time, indicating superiority of the tolerance mechanism of LPE at this concentration (Figure 2D).

We then examined the genetic changes in HPE lineages. Interestingly, none of the strains evolved at higher paraquat concentration showed polyamine transporter related mutations. Rather, all three lineages of HPE acquired an insertion element, IS1, upstream of the *emrE* gene (Figure 3A). Since such insertions can impact the expression levels ^10,23,24^, we examined the expression of *emrE* in evolved strains and WT. We observed a significantly higher expression of *emrE* in HPE (Figure 3B). The gene *emrE* encodes for a multidrug efflux pump that is implicated in paraquat export ^25,26^. The deletion of *emrE* in HPE resulted in complete loss of paraquat tolerance (Figure 3C). These observations suggest functionally opposite flux control strategies for paraquat tolerance and growth optimization in HPE as compared to LPE.

**Figure 3:**
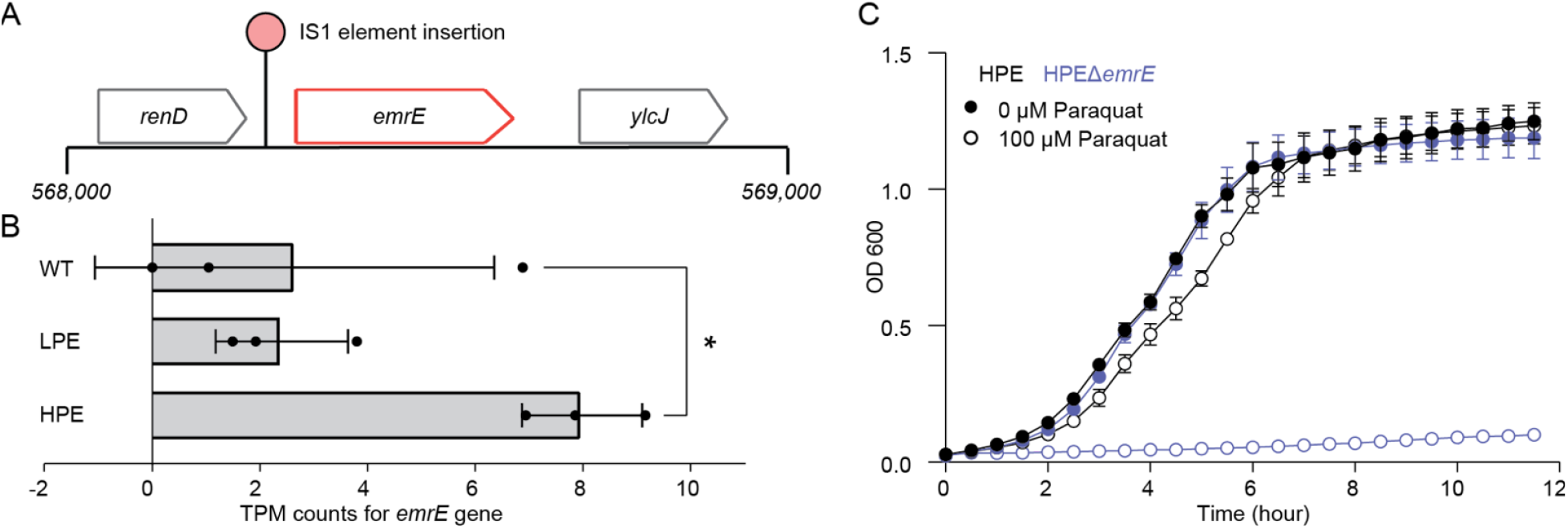
Adaptive strategy of oxidative stress tolerance in high-paraquat evolved *E. coli*. (A) Schematic for the mutation observed in the HPE strains. (B) Expression of the *emrE* gene in LPE and HPE strains. All three individual biological replicates are shown on the plot. Error bars represent standard deviation, and significance was determined using the Kruskal-Wallis test (*: P-value = 0.0253). (C) Effect of *emrE* gene deletion on the growth profile of HPE strain in the presence of 100 μM paraquat. The growth curve shows a mean of three biological replicates (with three technical replicates each), and the error bars show the standard error of the mean.

### Metabolic influence of different tolerance strategies

The specialization in the tolerance strategies motivated us to examine physiological logic behind the distinct evolutionary preference. We started with examining the expression status of the genes encoding canonical ROS detoxifying enzymes (Figure 4A). WT showed an expected paraquat dose-dependent upregulation of superoxide dismutases and other enzymes. LPE strains did not upregulate these enzymes at 10 μM paraquat concentration likely due to reduced influx of paraquat and hence lesser stress. However, at higher paraquat concentration LPE is required to upregulate ROS mitigating enzymes suggesting limited flux-restriction based tolerance. In contrast, HPE strains showed higher expression of these enzymes and are likely using both enzymatic detoxification and elevated paraquat efflux to maintain redox homeostasis.

**Figure 4:**
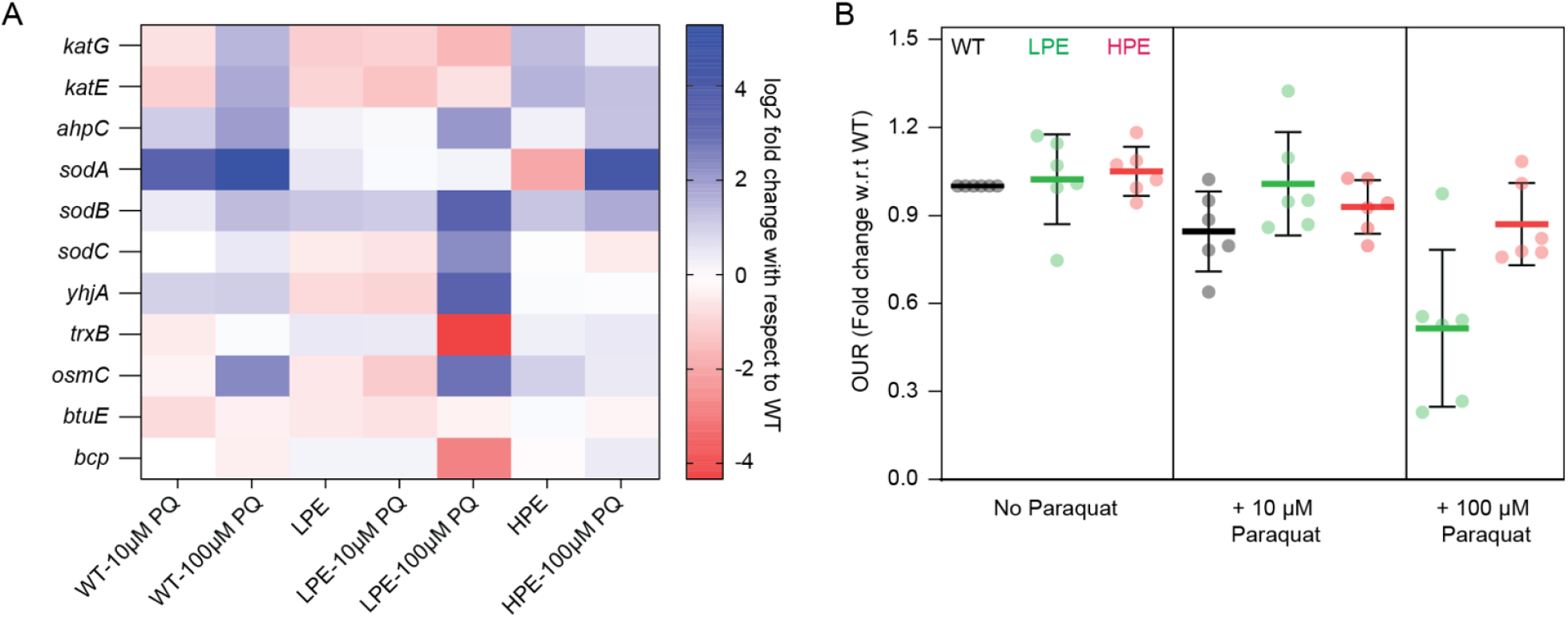
Transcriptional and respiratory responses to paraquat exposure. (A) Heatmap showing expression levels (log2 fold change with respect to WT) of the ROS mitigation related genes when treated with paraquat (PQ). The data is an average of three biological replicates. (B) Oxygen uptake rates of the strains in presence and absence of the specified paraquat concentrations. All six individual biological replicates shown on the plot. Error bars represent standard deviation. Significance was determined using the Kruskal-Wallis test and except for LPE-100 μM Paraquat (which showed ** significance), all others showed non-significance in comparison to WT-No Paraquat. (**: P-value = 0.0032)

Aerobic respiration is the primary source of ROS, and paraquat stress is reported to move *E. coli* to lower aerotype wherein energy dependency on ETS is reduced ^10,27^. We estimated the oxygen uptake rates of strains with various paraquat concentrations (Figure 4B). We did observe a reduction in oxygen consumption in LPE at higher levels of paraquat, a potential strategy to reduce ROS generation. However, the higher paraquat tolerance of HPE strains allowed them to avoid this metabolic downshift. Thus, the LPE and HPE strains seem to be engaging two different metabolic programs to deal with their respective redox stress.

### Cost associated with deploying different tolerance strategies

Our findings show that LPE partially decreases paraquat uptake, whereas HPE enhances its export alongside increased ROS detoxification. Although both approaches successfully alleviate redox stress, the reason for the dominance of one strategy over the other under conditions of persistently low versus high stress is not clear. To investigate this, we employed a computational analysis to determine the relative costs of each distinct strategy, motivated by the principle that bacterial metabolic choices are often governed by resource allocation constraints ^28,29^.

For this purpose, we used the StressME model ^30^, a unified genome-scale model of *Escherichia coli* metabolism, gene expression, and stress responses that can compute complete metabolic network rewiring under various stresses. The model accounts for the cellular costs of transports: the proton motive force (PMF) consumed by EmrE and the ATP used to operate PotD-PotABC. Consequently, the energy consumption (in the form of PMF or ATP) of both transporters can be simulated according to the adaptation strategy (LPE vs. HPE).

We incorporated the paraquat transport and redox-cycling pathway into StressME by curating and updating it with four coupled reactions representing paraquat import, intracellular redox cycling, ROS generation, and export (Figure 5A; Materials and Methods). Specifically, reactions 1 and 4 represent paraquat influx and efflux, respectively, while reactions 2 and 3 capture paraquat reduction and re-oxidation, which together generate superoxide radicals. This explicit representation enables simultaneous accounting of energetic costs, protein investment, and redox consequences associated with alternative detoxification strategies. Using the updated model, we quantified PMF and ATP demands associated with paraquat handling under strain-specific optimal split ratios and extracellular paraquat concentrations, enabling direct comparison of the energetic burden imposed by the adaptive strategies of the LPE and HPE strains.

**Figure 5:**
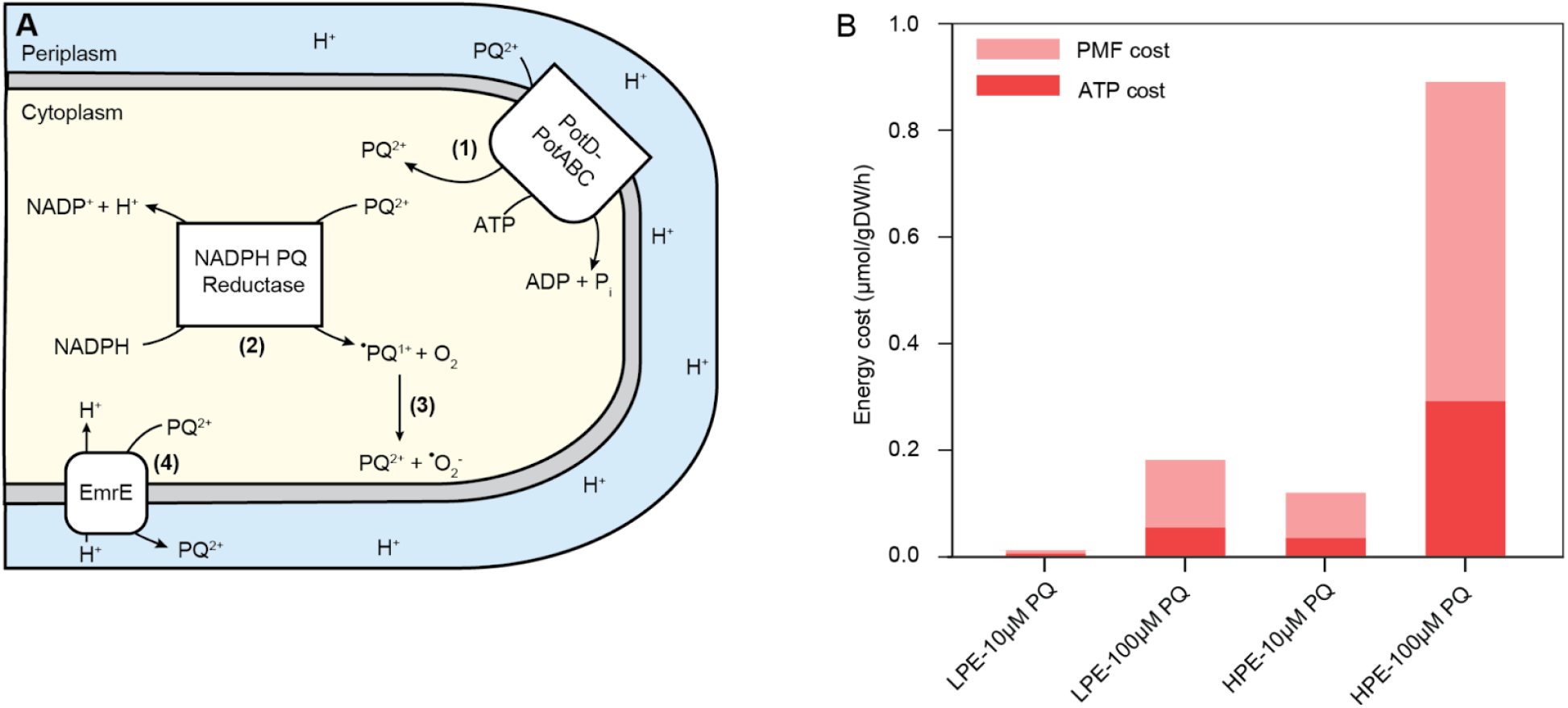
Energetic costs and fitness trade-offs associated with magnitude of paraquat stress. (A) Reactions of paraquat transport and redox cycling pathway that were implemented in a modified StressME model. (B) Genome-scale model based theoretical energy cost (PMF and ATP) estimates of the paraquat evolved strains when exposed to 10μM and 100μM paraquat.

The simulation showed a difference in PMF and ATP requirements between LPE and HPE strains. Both at 10 μM and 100 μM paraquat concentrations, LPE invested lesser energy resources as compared to HPE (Figure 5B). These cost estimations suggest that, while effective at high stress, the genetically fixed, high-energy adaptive approach of HPE is disadvantageous at lower stress levels. This difference in energy demand further explains the observation of LPE outcompeting HPE at 10 μM paraquat concentration (Figure 2D) and the logic behind evolutionary selection of entry restriction based strategy for the lower paraquat evolution paradigm.

### Tolerance upshift in evolved strains

The two distinct adaptive mechanisms specific for different ranges of paraquat motivated us to delineate additional layers of tolerance strategies in *E. coli*. Towards this objective, we designed a series of ALE experiments (Figure 6A). To elevate the stress tolerance, we evolved LPE strain under 100 μM paraquat stress. All three lineages of the LPE strain evolved to 100 μM paraquat (designated as LH1PE) acquired earlier described efflux strategy, similar to HPE strains, to improve their tolerance (Supp. Table 2). However, LH1PE strain when further evolved at doubled paraquat concentration (200 μM; designated as LH2PE) fixed mutations in the gene encoding the folate-dependent protein YgfZ.

**Figure 6:**
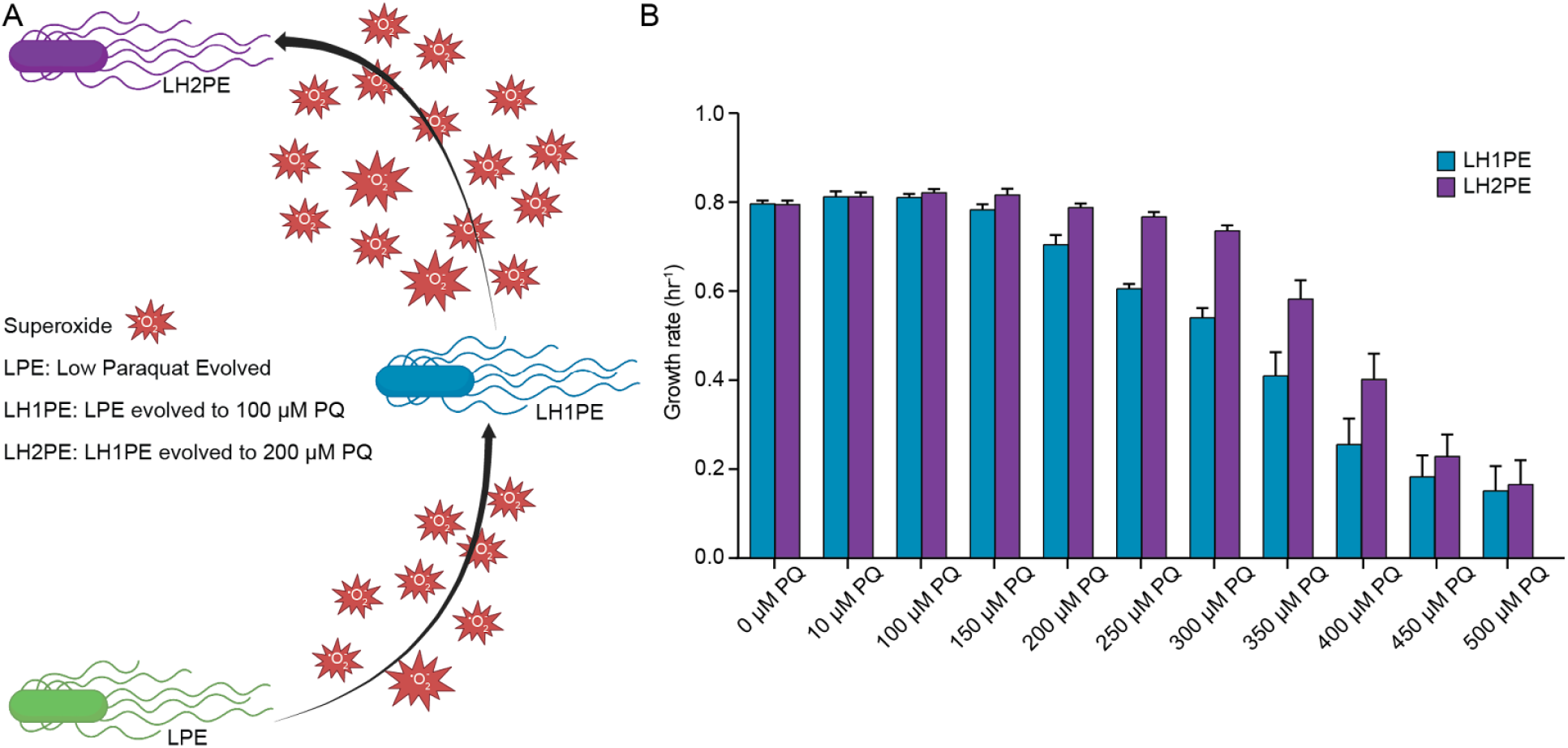
Adaptive evolution of LPE in presence of higher levels of superoxide. (A) Schematic of the adaptive laboratory evolution (ALE) design. (B) Growth rates of the evolved strains upon exposure to different paraquat (PQ) concentrations. The plot shows the mean of three biological replicates (each with three technical replicates), and the error bars show the standard error of mean.

The mitochondrial homolog of YgfZ, IBA57, is involved in the assembly of the Fe-S cluster ^31^. Though poorly characterized, YgfZ is believed to function similarly to its homolog and has been implicated in oxidative stress mitigation ^32^. We therefore examined whether the mutational targeting of *ygfZ* is directed to achieve additional tolerance. We performed a comparative sensitivity profiling of LH1PE and LH2PE using a range of paraquat concentrations (Figure 6B). Both strains showed similar tolerance till 100 μM paraquat, but beyond this concentration LH2PE started to show a clear dose-dependent tolerance superiority. This observation highlights the metabolic augmentation of ROS defense built upon flux control strategies.

### Conclusion

Collectively, our results establish that bacterial oxidative stress tolerance is organized as a hierarchical and resource-aware decision process rather than a uniform amplification of defense pathways. By demonstrating that *E. coli* deploys distinct, stress-magnitude-specific modules - ranging from low-cost restriction of toxic influx to energetically intensive efflux and detoxification programs - we uncover a general principle by which cells balance robustness against metabolic economy. This operating cost distinction explains why certain adaptive solutions are favored only within defined stress regimes and predicts when fixed, high-cost defenses become maladaptive. Moreover, the ability of flux-control strategies to serve as a foundation for subsequent metabolic rewiring reveals how incremental tolerance gain can emerge without wholesale regulatory redesign. These insights extend beyond oxidative stress, offering a conceptual framework for understanding how microbes evolve graded, context-dependent responses to diverse environmental challenges, with implications for infection biology, antibiotic resistance evolution, and the rational design of stress-targeting therapies.

## Materials and Methods

### Bacterial strains and growth conditions

*Escherichia coli* K12 MG1655 (ATCC 700926) was used as the wild-type strain. All the growth assays were performed using the Tecan Spark multimode microplate reader at 37°C and atmospheric oxygen partial pressure using 96 well plate with 200μL culture per well. At least three biological replicates (each with three technical replicates) were performed for each experiment. Unless stated otherwise, M9 minimal medium supplemented with 4 g/L glucose was used for bacterial growth. For the growth profiling with media variation, paraquat was added to M9 minimal medium supplemented with 4 g/L glucose at the required concentrations.

*potD, potF* and *emrE* knockout strains were generated using the P1 phage transduction method, with Keio collection strains as donors for gene knockout cassettes ^33,34^.

Growth rate calculations were done using gcplyr: an R package for microbial growth curve data analysis ^35^.

### Adaptive laboratory evolution

Adaptive laboratory evolution was performed using three independent replicates of the *E. coli* wild-type strain. Low paraquat evolution was performed by serially propagating the cultures in M9 minimal medium supplemented with 4 g/L glucose and 10 μM paraquat at 37°C and atmospheric oxygen partial pressure, while the high paraquat evolution was performed in the same conditions with 100 μM paraquat. Cultures were always maintained in excess nutrient conditions assessed by non-tapering exponential growth. The laboratory evolution was performed for a sufficient interval to allow the cells to reach their growth rate plateau.

Further evolution of LPE at 100 μM paraquat and LH1PE at 200 μM paraquat was performed in a similar manner.

### DNA resequencing

DNA resequencing was performed on clones from the endpoints of evolved strains. Total DNA was sampled from an overnight culture and immediately centrifuged for 5 min at 12,000 r.p.m. The supernatant was decanted, and the cell pellet was frozen at −80°C. Genomic DNA was isolated using QIAamp PowerFecal Pro DNA Kit (Cat No./ID: 51804, QIAGEN GmbH) following the manufacturer’s protocol. Resequencing libraries were prepared using QIAseq® FX DNA library kit from Qiagen following the manufacturer’s protocol. Libraries were run on a HiSeq and/or NextSeq (Illumina).

Sequencing reads were filtered and trimmed using the FastP software. The breseq bioinformatics pipeline version 0.31.1 was used to map sequencing reads and identify mutations relative to an *E. coli* K-12 MG1655 reference genome (NC_000913.3) amended to reflect the starting strain best.

### Structure rendering and visualization in PyMol

Protein structure of PotD-PotABC (PDB ID: 8Y5I) of *E. coli* K-12 was acquired from the Protein DataBank (PDB). The structure was rendered using PyMol to facilitate detailed visualization.

### RNA sequencing

Bacteria were grown in M9 minimal medium supplemented with 4 g/L glucose, with respective paraquat concentration, till their mid-exponential phase. The culture was harvested and immediately processed for RNA isolation. The cells were dissolved in TRIzol and incubated at 65°C for 15 min. RNA was obtained in an aqueous layer upon the addition of chloroform, which was then precipitated using chilled isopropanol. This RNA pellet was washed with alcohol and finally resuspended in RNase-free water.

Total RNA concentration was measured using the Qubit RNA High Sensitivity assay according to the manufacturer’s instructions, and RNA integrity and fragment size distribution were evaluated using the Agilent Fragment Analyzer with a high-sensitivity RNA kit. RNA sequencing libraries were prepared using the QIAseq FastSelect rRNA depletion system (5S/16S/23S) in combination with the QIAseq RNA library preparation enzymes and buffers, following the manufacturer’s protocols. After library construction, final library concentrations were quantified using the Qubit dsDNA High Sensitivity assay, and library size distribution and quality were assessed with the Agilent high-sensitivity NGS fragment analysis system. Libraries were normalized and pooled at equimolar concentrations prior to sequencing, and the pooled libraries were sequenced on an Illumina NextSeq 2000 platform. Further analysis was performed as described in a previous study ^36^.

### Phenotype characterization

Culture density was measured at 600 nm absorbance with a spectrophotometer and correlated to cell biomass. Samples for exometabolite estimation were filtered through a 0.22 μm filter (PVDF, Millipore) and measured using refractive index detection by HPLC (Agilent 12600 Infinity) with a Bio-Rad Aminex HPX87-H ion exclusion column. The HPLC method was the following: injection volume of 10 μL and 5 mM H_2_SO_4_ mobile phase set to a flow rate and temperature of 0.5 mL/min and 45°C, respectively, for a run time of 20 min per sample.

The oxygen uptake of each culture was determined by measuring the depletion of dissolved oxygen using the Oroboros O2K system. The cultures were grown in M9 minimal medium supplemented with 4 g/L glucose, with respective paraquat concentration, at 37°C and atmospheric oxygen partial pressure, and oxygen uptake rates were normalized to OD600.

### Competition assay and relative fitness estimation

The relative fitness of paraquat-tolerant strains was assessed using pairwise competition assays. Log-phase monocultures of the WTΔ*potD* strain (representative of the LPE strain) and the HPE strain (lineage A) were mixed in a 1:1 ratio and inoculated into 15 mL of M9 minimal medium supplemented with 4 g/L glucose and 10 μM paraquat.

The mixed culture was incubated at 37°C and atmospheric oxygen partial pressure to allow competition between the strains. Samples were collected at three defined time points: early (0.5 h), mid (2 h), and late (7 h). At each time point, cultures were serially diluted and plated onto Luria-Bertani (LB) agar plates to determine total viable counts.

To distinguish WTΔ*potD* from the competing population, individual colonies from LB plates were patch-plated onto LB agar containing kanamycin. Colonies that grew on LB-kanamycin plates were scored as WTΔ*potD*. The competitive index was calculated as the proportion of WTΔ*potD* colonies within the total population using the following formula:

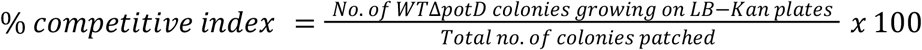

### Simulation of Paraquat Transport and Redox Cycling

The paraquat (pq2) transport and redox cycling pathway was implemented in a modified StressME model (Docker version: queensysbio/stressme:v1.2) through four coupled reactions (Eqs. 1 - 4). Equation (1) represents the PotD-PotABC-mediated import of pq2 from the periplasm to the cytoplasm (*v*_import). Equation (2) describes the enzymatic reduction of cytoplasmic pq2 to pq1 by PQ2RED, encoded by *b3924* (*v*_reduction). Equation (3) represents the oxidation of pq1 back to pq2 by PQ1OX under aerobic conditions (*v*_oxidation), during which molecular oxygen serves as the electron acceptor and superoxide (^•^O_2_^−^) is produced as a byproduct. The generated superoxide contributes to cellular oxidative stress, leading to protein damage and activation of detoxification pathways (e.g., superoxide dismutase and catalase systems), which may impose additional protein synthesis and energy costs. Finally, Equation (4) describes pq2 export to the periplasm via the multidrug efflux transporter EmrE, encoded by *b0543* (*v*_export).

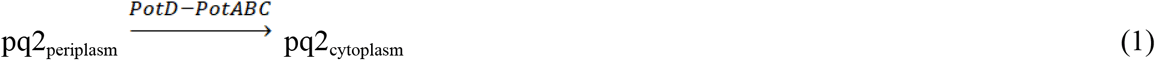

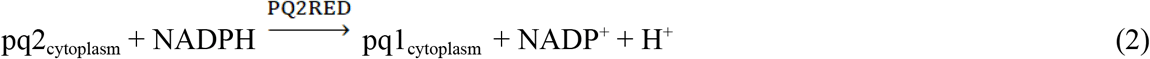

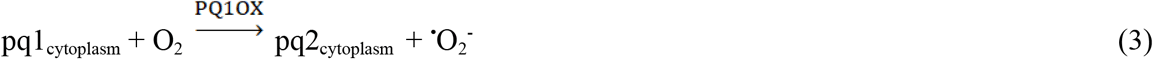

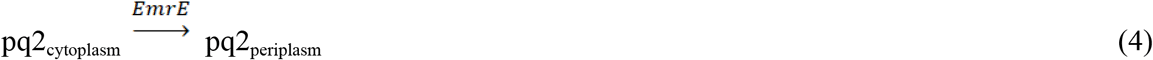

The relationship between extracellular pq2 concentration ([pq2]_ext_) and PotD-PotABC uptake flux (*v*_import) was described by a Michaelis-Menten function (Eq. 5):

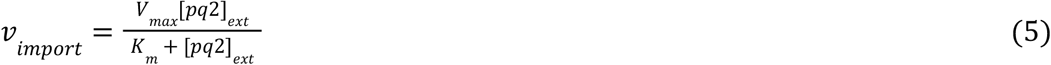

where V_max_ and K_m_ were calibrated against experimental paraquat uptake data vs. growth rates. Optional competitive inhibition can be modeled by including inhibitor and inhibition constant (K_i_) terms, modifying K_m_ as K_m_^eff^ = K_m_(1+[*I*]/K_i_).

Mass balance for pq2 in the cytoplasm was enforced as a constraint in the model:

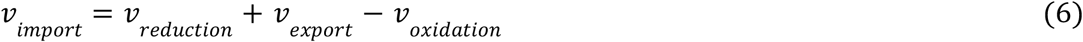

Under steady-state redox cycling, v_reduction_ = v_oxidation_, simplifying the relationship to v_export_=v_import_. The flux partition between reduction and export was defined by a flux split ratio (frac_red_), where v_reduction_ = frac_red_ x v_import_. The optimized values frac_red_ = 0.2 for the HPE strain and frac_red_ = 1.0 for the LPE strain were calibrated against the experimental data ([pq2]_ext_ vs. growth rates).

Protein abundance ratio between PQ2RED and EmrE ([P_red_]/[P_EmrE_]) was approximated from transcript counts measured by RNA-seq, assuming comparable translation efficiencies (TE≈1). Thus,

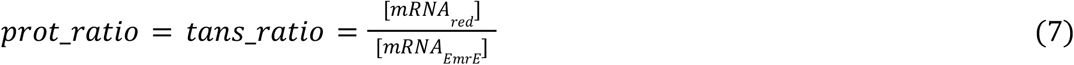

and the effective catalytic rate constant for EmrE was estimated as

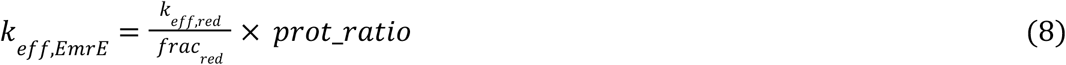

The computed fluxes of v_reduction_ and v_export_ = v_import_-v_reduction_+v_oxidation_ (Eqs. 6–8) were then used to set fixed upper and lower bounds for the corresponding reactions PQ2RED and EmrE within the StressME model. This formulation ensures consistent paraquat mass balance while linking experimental transcript and phenotype data to pq2 transport, enzyme turnover efficiency and stress-induced energy demand.

### Cost-Benefit Analysis of EmrE-Mediated Paraquat Export

To quantify the cellular cost and benefit of EmrE-mediated paraquat (pq2) export, we analyzed fluxes from the StressME model under an optimized split ratio (i.e., 0.2 and 1.0 for the HPE and LPE strain, respectively) using the simulated full flux dataset (Supplementary Information Dataset S1) under different extracellular paraquat concentrations, where the growth rate (μ, h^−1^) and translation fluxes for EmrE (t_EmrE_ representing translation_b0543) and a pq2 reductase (t_Red_ representing translation_b3924) were extracted (mmol protein synthesized gDW^−1^ h^−1^).

pq2 fluxes were partitioned into import (v_import_), intracellular reduction (v_reduction_), and export (v_export_) through EmrE (all in mmol gDW^−1^ h^−1^). The energetic costs associated with transport were calculated as:

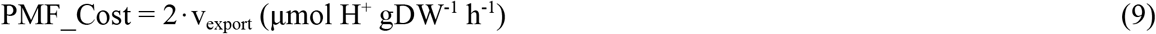

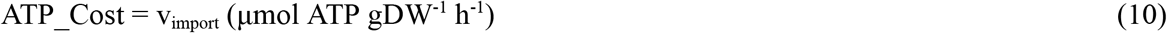

assuming two protons per pq2 exported and one ATP per pq2 imported. This framework enables direct assessment of the trade-off between energy and protein investment on pq2 export versus reduction via EmrE-mediated export and PQ2RED-mediated reduction, respectively.

## Data Availability

DNAseq and RNAseq data supporting this study are deposited in the NCBI SRA (PRJNA1404458) and GEO (GSE316857), respectively. The computational environment used for all simulations is available as a Docker container (stressme v1.2) from Docker Hub at https://hub.docker.com/r/queensysbio/stressme/tags (Tag: v1.2; Digest: sha256:b41b5348235dcd4cb37c608e89cca3e53d72ecb67bd29c5b3bf491f099e81d12). The flux solutions generated from the Low Paraquat Evolved (LPE) and High Paraquat Evolved (HPE) simulations are included in the Supplementary Information Dataset S1.

## Acknowledgement

This work was funded by the DAE, India-Tata Institute of Fundamental Research Grant and Ramalingaswami Re-entry Fellowship, DBT, India to A.A. This work also received funding by the Government of Canada through Genome Canada and Ontario 904 Genomics (OGI-207), and Genome Quebec to L.Y. We acknowledge the support of NGS Facility at NII, India. We thank Neha Banwani for her help with some validation experiments for the project. Icons for some figures were created using our license at Biorender.com.

## Author contributions

A.A. and A.P. designed the study. A.P., S.K. and S.S. performed experiments. A.P., S.K., and A.A. analyzed the data. J.Z. and L.Y. performed the computational analyses. A.P., and A.A. wrote the manuscript with contributions from all other co-authors.

## Supplementary Table

**Supp. Table 1:** Sample code used during running the ALE experiment.

| Strain | ALE code |
| --- | --- |
| LPE-A | *E. coli*_A9F60I1 |
| LPE-B | *E. coli*_A10F60I1 |
| LPE-C | *E. coli*_A12F60I1 |
| HPE-A | *E. coli*_A25F45I1 |
| HPE-B | *E. coli*_A26F45I1 |
| HPE-C | *E. coli*_A27F45I1 |
| LH1PE-A | *E. coli*_A29F40I1 |
| LH1PE-B | *E. coli*_A30F40I1 |
| LH1PE-C | *E. coli*_A31F40I1 |
| LH2PE-A | *E. coli*_A37F30I1 |
| LH2PE-B | *E. coli*_A38F30I1 |
| LH2PE-C | *E. coli*_A39F30I1 |

**Supp. Table 2:** List of genes mutated in the paraquat evolved strains.

| Strain | Mutations compared to WT *E. coli* strain |  |  |  |
| --- | --- | --- | --- | --- |
| LPE-A | *rpoB*<br>Y555H | *potC*<br>IS2 Coding (497-501/795 nt) |  |  |
| LPE-B | *rpoB*<br>H526Y | *potD*<br>Δ1bp Coding (260/1047 nt) |  |  |
| LPE-C | Δ82bp<br>*pyrE*→/→*rph* | *potA*<br>G55D |  |  |
| HPE-A | *rpoB*<br>S712C |  | *renD*→/→*emrE*<br>IS1 Intergenic<br>(+21/-40) |  |
| HPE-B | *rpoB*<br>S712C |  | *renD*→/→*emrE*<br>IS1 Intergenic<br>(+21/-40) |  |
| HPE-C | *rpoB*<br>H526Y |  | *renD*→/→*emrE*<br>IS1 Intergenic<br>(+21/-39) |  |
| LH1PE-A | *rpoB*<br>H526Y | *potD*<br>Δ1bp Coding (260/1047 nt) | *renD*→/→*emrE*<br>IS1 Intergenic<br>(+21/-39) |  |
| LH1PE-B | *rpoB*<br>H526Y | *potD*<br>Δ1bp Coding (260/1047 nt) | *renD*→/→*emrE*<br>IS1 Intergenic<br>(+21/-39) |  |
| LH1PE-C | *rpoB*<br>H526Y | *potD*<br>Δ1bp Coding (260/1047 nt) | *renD*→/→*emrE*<br>IS1 Intergenic<br>(+21/-39) |  |
| LH2PE-A | *rpoB*<br>H526Y | *potD*<br>Δ1bp Coding (260/1047 nt) | *renD*→/→*emrE*<br>IS1 Intergenic<br>(+21/-39) | *ygfZ*<br>A65V |
| LH2PE-B | *rpoB*<br>H526Y | *potD*<br>Δ1bp Coding (260/1047 nt) | *renD*→/→*emrE*<br>IS1 Intergenic<br>(+21/-39) | *sdhE*←/→*ygfZ*<br>Intergenic (-232/-11) |
| LH2PE-C | *rpoB*<br>H526Y | *potD*<br>Δ1bp Coding (260/1047 nt) | *renD*→/→*emrE*<br>IS1 Intergenic<br>(+21/-39) | *ygfZ*<br>N215D |

## Supplementary Figure

**Supp. Figure 1:**
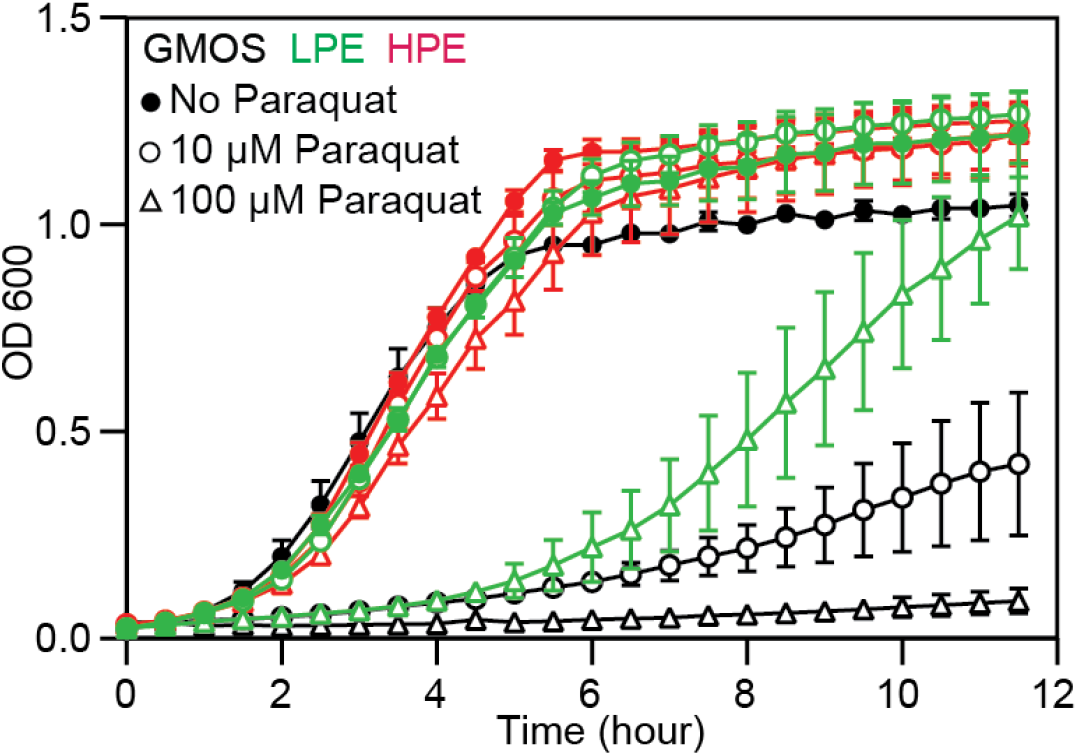
Impact of paraquat on the growth of GMOS, LPE and HPE strains. The growth curve shows a mean of three biological replicates (with three technical replicates each), and the error bars show the standard error of the mean.

